# R2Dtool: Integration and visualization of isoform-resolved RNA features

**DOI:** 10.1101/2022.09.23.509222

**Authors:** Aditya J. Sethi, Pablo Acera Mateos, Rippei Hayashi, Nikolay Shirokikh, Eduardo Eyras

## Abstract

Long-read RNA sequencing enables the mapping of RNA modifications, structures, and protein-interaction sites at the resolution of individual transcript isoforms. To understand the functions of these RNA features, it is critical to analyze them in the context of transcriptomic and genomic annotations, such as open reading frames and splice junctions. To enable this, we have developed R2Dtool, a bioinformatics tool that integrates transcript-mapped information with transcript and genome annotations, allowing for the isoform-resolved analytics and graphical representation of RNA features in their genomic context. We illustrate R2Dtool’s capability to integrate and expedite RNA feature analysis using epitranscriptomics data. R2Dtool facilitates the comprehensive analysis and interpretation of alternative transcript isoforms.

R2Dtool is freely available under the MIT license at: https://github.com/comprna/R2Dtool.

## Introduction

Long-read sequencing enables accurate mapping of diverse RNA features with transcript isoform resolution. Such features include, among others, chemical modifications (Acera Mateos et al., 2024; Bansal et al., 2024; Liu et al., 2019; Stephenson et al., 2020), structured regions (Aw et al., 2021; Stephenson et al., 2022), and RNA-protein interacting sites (Lin et al., 2022). The functional roles of these RNA features are often linked to their position relative to transcript landmarks, such as transcription start sites, splice junctions, start and stop codons, and polyadenylation sites. For example, positional enrichment of N6-methyladenosine (m6A) downstream of stop codons led to the discovery of an m6A role in transcript deadenylation at the 3’ end of transcripts (Dominissini et al., 2012; Lee et al., 2020). Similarly, the identification of isoform-specific exon secondary structures made possible a link with transcript-specific translational efficiency (Aw et al., 2021); and the positional distribution of RNA-protein interaction sites along transcripts revealed specific modes of RNA regulation (Lin et al., 2022; Van Nostrand et al., 2020). The isoform-resolved visualization of RNA features in the context of transcriptomic and genomic landmarks is thus critical to facilitate the discovery of isoform-specific regulatory mechanisms.

The transcriptome-wide distributions of RNA features are often represented using metagene plots, which describe the positional density of these features in the context of simplified gene models. However, current methods for generating metagene plots generally select a single representative isoform, often the longest one or most abundant, against which the positions of all RNA features are calculated (Fournier et al, 2019.; Olarerin-George and Jaffrey, 2017). This may result in a misassignment of distances and distributions of the RNA features relative to the transcriptomic or genomic landmarks at loci that simultaneously express multiple isoforms, which in mammals is expected to occur in around one in every five genes (Tapial et al., 2017). Isoform-resolved transcriptomic methods, such as long-read sequencing, offer a solution to this challenge by enabling the direct assignment of RNA features to specific transcripts. However, available metagene methods do not capitalize on this data to enable isoform-aware analysis.

Here, we describe R2Dtool, a computational method for the integration and visualization of isoform-resolved RNA features in the context of transcriptomic and genomic annotations. R2Dtool exploits the isoform-resolved mapping of RNA features, such as those obtained from long-read sequencing, to enable simple, reproducible, and lossless integration, annotation, and visualization of isoform-specific RNA features. We illustrate R2Dtool’s capabilities with the analysis of isoform-resolved messenger RNA (mRNA) modification data, which can be accurately obtained from nanopore long-read sequencing data (Acera Mateos et al., 2024; Hendra et al., 2022) and its interpretation in the right mRNA isoform context has uncovered important properties and regulatory mechanisms (Gleeson et al., 2024; Uzonyi et al., 2022).

## Implementation

R2Dtool starts with a set of RNA feature positions in transcriptomic coordinates, such as those obtained from long-read sequencing reads mapped directly to transcript isoforms (Acera Mateos et al., 2024; Gleeson et al., 2024; Hendra et al., 2022; Stephenson et al., 2022) (Fig. 1A). R2Dtool operates with these transcript features and the corresponding gene annotations to analyze, integrate, and visualize isoform-resolved RNA feature maps. R2Dtool’s core function *liftover* transposes the transcript-centric coordinates of the isoform-mapped sites to genome-centric coordinates. Another core function, *annotation*, performs positional annotation of RNA features with isoform-aware metagene coordinates and distances to annotated transcriptomic and genomic landmarks. Furthermore, R2Dtool provides isoform-aware visualization in the form of *metaplots*. These include the *metatranscript* plot, which visualizes the isoform-specific distribution of RNA features in the context of a rescaled mRNA coordinate system. Additionally, R2Dtool enables visualization of the positional distribution of RNA features around transcript landmarks, such as isoform-specific splice junctions (*metajunction* plot) and start or stop codons (*metacodon* plot).

**Figure 1.**
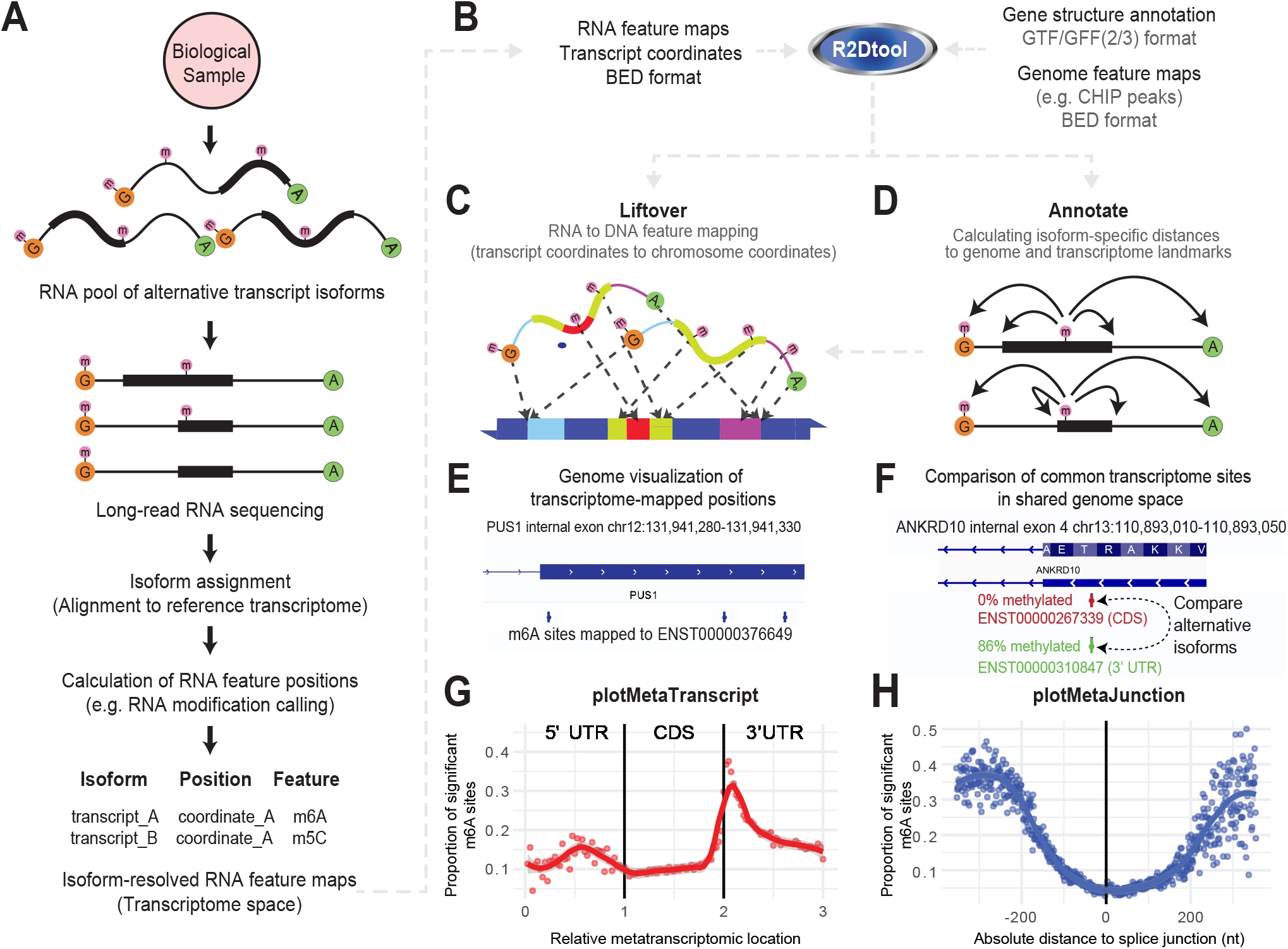
R2Dtool facilitates the analysis and interpretation of RNA isoform-specific features in the context of transcriptomic and genomic annotations. **(A)** Diagram describing the analysis steps of long-read RNA profiling experiments resulting in the generation of RNA feature maps in transcript-centric coordinates. **(B)** Inputs, outputs, and operations of R2Dtool. **(C)** Schematic for R2Dtool *liftover* functionality, where isoform-resolved sites are annotated with their genomic coordinates, using a GTF file to map between the two coordinate systems. **(D)** Schematic of R2Dtool *annotate* functionality, which calculates isoform-specific distances between RNA feature positions and RNA landmarks which are annotated on the same isoform. **(E)** R2Dtool liftover enables visualization of transcriptome-mapped m6A sites in their genomic context. IGV snapshot shows three m6A sites mapped to transcript ENST00000376649 in the context of PUS1 gene architecture. **(F)** IGV browser view of transcript-specific m6A sites in two different ANKRD10 isoforms with different stoichiometry that share identical genomic positions. The longer isoform (red) shows 0% methylation at the highlighted site, where the site lies in the transcript CDS. The shorter transcript (green) shows 86% m6A stoichiometry at the same genomic position, which lies in the transcript 3’ UTR. **(G)** The plot shows the output of R2Dtool *plotMetaTranscript*. The x-axis shows R2Dtool *annotate* metatranscript coordinates. Each point represents the average modification rate in a given metatranscript bin, 5’ untranslated region (UTR), open reading frame (ORF), and 3’UTR of human mRNA transcripts, where the average modification rate is calculated as the number of modified sites over the number of tested sites in the bin. A LOESS trendline is shown across the data, with the shaded area representing the 95% confidence interval for the LOESS fit. (H) The plot shows the output of R2Dtool *plotMetaJunction*, revealing the density of m6A modifications relative to the distances to the closest upstream (left) and downstream (right) exon-exon junction in all the mRNA transcript isoforms to which m6A sites were mapped. Each point represents the average modification rate at a given 1nt interval from the nearest splice junction, where the average rate is calculated as the number of modified sites over the number of tested sites in the 1nt interval. As in G, a LOESS trendline is shown across the data, with the shaded area representing the 95% confidence interval for the LOESS fit. For G and H, significant sites were defined as those predicted by Dorado with >10% m6A stoichiometry, calculated by ModKit.

R2Dtool is applicable to any organism with a transcriptome annotation, either obtained from an existing reference or user built. R2Dtool’s functions are implemented as standalone command-line tools to follow pipelining and format preservation principles. This facilitates rapid and flexible coupling with other bioinformatic workflows, such as RNA modification profiling, isoform-resolved RNA structure detection, or protein interaction studies. R2Dtool can also be applied to short-read experiments in cases where RNA features can be mapped to specific isoforms, e.g. when analyzing *in vitro* transcribed sequences that map to specific transcripts. Software and usage details are provided at https://github.com/comprna/R2DTool.

### Input and output formats

R2Dtool is designed to work with RNA features along transcripts described in BED3+ format (Niu et al., 2022) and gene annotations provided in gene transfer format (GTF2) (Pertea and Pertea, 2020) (Fig. 1B). Columns 1-3 of the input BED3+ file must provide the positions of the RNA features on the reference transcriptome. This BED3+ input file can contain any number of additional columns to encode experiment-specific data, including scores, labels, likelihoods, or any metadata generated during the upstream analysis, in concordance with the BED3+ definition. R2Dtool uses gene information from a GTF2 input file, which minimally includes the annotations of coding and non-coding exons but can incorporate additional information. This enables the integration and comparison of the transcript-resolved data with genomic annotations (Fig. 1B-D). R2Dtool generates BED6+n files as outputs including the transcriptomic and genomic coordinates of given sites, together with additional positional properties. The BED6+ output file format is compatible with other tools that handle BED files in genome-based coordinates, such as Samtools, Bedtools, and the IGV genome browser (Li et al., 2009; Quinlan, 2014; Thorvaldsdottir et al., 2013).

### Liftover of RNA-mapped features to DNA coordinates

To enable the comparison between isoform-mapped RNA features and genomic annotations, R2Dtool lifts the positions of transcriptome-mapped RNA features over to the corresponding genome reference, using a GTF2 transcript annotation file to calculate the relevant genome positions (Fig. 1C). This operation is performed with the command ‘`r2d liftover -i <Sites (BED3+)> -g <Annotation (GTF2)>`‘. The output is a BED6+ file, where columns 1-6 contain the new genomic coordinates of each RNA feature in standard BED6 format, whereas the input data, including the original transcriptome-reference coordinates, are losslessly preserved in the output columns 7:n+7, where *n* was the original number of columns in the input data. To illustrate this operation, we used R2Dtool’s *liftover* to visualize m6A modification calls generated from nanopore direct RNA sequencing of HeLa transcriptomic RNA (SQK-RNA004 kit). DRACH-context m6A basecalling was performed with Dorado v0.5.3 (https://github.com/nanoporetech/dorado) and the methylation calls were processed with ModKit v0.2.6 (https://github.com/nanoporetech/modkit), before analysis by R2Dtool (code for this analysis is available at https://github.com/comprna/R2Dtool/). While the m6A calls were made in transcript-centric coordinates, R2Dtool’s *liftover* enabled the visualization of m6A sites in their genomic context. This is illustrated by showing different m6A sites in a transcript from gene PUS1 in their genomic context and in proximity to an adjacent intron (Fig. 1E). This operation highlights that by transposing sites from alternative transcript isoforms to their genomic coordinates, it is possible to assess features in their right molecular context that would otherwise be missed by genomic-based methods.

### Isoform-specific positional annotation

R2Dtool leverages the specificity of isoform-resolved RNA feature maps to systematically annotate transcript-mapped RNA feature positions with absolute and relative distances to transcriptomic and genomic landmarks, including transcript starts and ends, splice junctions, stop codons and start codons (Fig. 1D). This operation is performed with the command ‘`r2d annotate -i <Sites (BED3+)> -g <Annotation (GTF2)>`‘, where the first 3 columns of the input specify the position of the features in transcriptome-specific BED3 coordinates, assumed to be on the plus strand. The output of this command is a BEDn+12 file, where the original *n* columns of the input are preserved, and 12 additional columns are added to the output. These additional columns correspond to the *gene ID, gene name, transcript biotype*, feature metatranscript coordinates, and the absolute and relative distances to local features, such as stop and start codons and adjacent splice junctions.

### Isoform-aware metatranscript coordinates and plots

R2Dtool calculates an isoform-aware metacoordinate for features localized on transcripts that are annotated as protein-coding. This metacoordinate represents the normalized position of a given feature on a fixed-length virtual transcript model using the same method as previously described, where the positions of features are linearly scaled to a position between 0-3, depending on the relative position of the feature across the span of the 5’ UTR [0-1), CDS (1-2), or 3’ UTR (2-3] (Olarerin-George and Jaffrey, 2017). However, unlike other metagene methods that use a single transcript model to calculate the metacoordinates for all features assigned to a given gene, R2Dtool’s leverages the individual isoform assignment of each RNA feature to calculate distinct metacoordinates for each alternative transcript that spans a given genomic position, accommodating potential diversity in UTR and ORF start/end positions present in alternative transcript isoforms. We highlight this capability by showing how a single genomic position on ANKRD10 is heavily m6A methylated when mapped to a transcript isoform where the site is located in the isoform 3’ UTR (86% m6A/A; metacoordinate of 2.26), but unmethylated when mapped to an alternative isoform, where the same genomic position corresponds to the transcript CDS (0% m6A/A, metacoordinate of 1.54) (Fig. 1F).

Metatranscripts coordinates are calculated for all features mapped onto protein-coding RNAs during the *annotation* analysis step and can be readily visualized through publication-grade isoform-resolved feature distribution plots using R2Dtool’s plotting functions. These include the metatranscript plot, which shows the normalized positional density of RNA features. The R2Dtool command ‘`r2d plotMetaTranscript’` produces density plots showing the distribution of RNA features from the output of the *annotate* command. The density of RNA features is determined across bins spanning the virtual metatrancript model by comparing the proportion of *positive* RNA features (e.g. with stoichiometry above or *p*-value below a certain cutoff), compared to all tested sites in the given metatranscript bin. For the metatranscript plot, we segment the metatranscript into 120 bins of width 0.025. To illustrate this operation, we produced a metatranscript plot for the density of m6A sites with >10% stoichiometry along R2Dtool-annotated transcript-centric m6A calls performed in HeLa (Fig. 1G).

### Isoform-specific landmark-centric plots

R2Dtool also calculates the distribution of absolute distances between RNA features and reference landmarks, such as the start and end of the ORF, the start and end of the transcript, or the nearest upstream or downstream splice junction, all in an isoform-specific manner, enabling a range of downstream positional analyses. We illustrate this operation with the analysis of the relative distances of m6A modifications to their nearest upstream or downstream exon-exon junctions obtained from the *annotate* command above (Fig. 1H). This plot, produced with the command ‘`r2d plotMetaJunction`’, shows a clear exclusion of m6A sites in a window of 200nt around the exon-exon junctions, in agreement with previous reports (Uzonyi et al., 2022). Like the metatranscript plot, the metajunction plot displays the proportion of positive features at intervals around each splice-site, in increments of 1nt. Similarly, ‘`r2d plotMetaCodon`’ can be used to plot the distribution of RNA features in absolute distance around start and stop codons, also in 1nt intervals (also available at https://github.com/comprna/R2DTool).

## Conclusions

R2Dtool provides a simple, robust, and reproducible framework to integrate and visualize isoform-resolved RNA feature maps, enabling a comprehensive annotation of transcript-centric sites. R2Dtools provides isoform-aware identification of RNA feature distribution and can inform on potential RNA regulatory mechanisms from long-read transcriptomics. R2Dtool is particularly useful for epitranscriptomics, where there have been recent significant advances in the isoform-specific identification of RNA modifications using long-read direct RNA sequencing, where most methods generate their estimates in transcript-centric coordinates. We anticipate that R2Dtool could empower future epitranscriptomic studies to discover new RNA regulatory mechanisms, particularly those involving the interplay between RNA modifications and genomic features. By seamlessly integrating transcriptomic and genomic annotations using standard formats, R2Dtool empowers researchers to fully utilize the potential of isoform-resolved transcriptomics.

## Data availability

Data produced in this work is available from https://doi.org/10.6084/m9.figshare.25730082.v1

